# Anti-CRISPR phages cooperate to overcome CRISPR-Cas immunity

**DOI:** 10.1101/279026

**Authors:** Mariann Landsberger, Sylvain Gandon, Sean Meaden, Hélène Chabas, Angus Buckling, Edze R. Westra, Stineke van Houte

**Affiliations:** ESI and CEC, Biosciences, University of Exeter, Cornwall Campus, Penryn TR10 9EZ, UK; CEFE UMR 5175, CNRS Université de Montpellier Université Paul-Valéry Montpellier EPHE, 34293 Montpellier Cedex 5, France

**Keywords:** CRISPR-Cas, bacteria, phage, partial resistance, immunosuppression, anti-CRISPR, epidemiology, tipping points, bifurcation, Allee effect

## Abstract

Some phages encode anti-CRISPR (*acr*) genes, which antagonize bacterial CRISPR-Cas immune systems by binding components of its machinery, but it is less clear how deployment of these *acr* genes impacts phage replication and epidemiology. Here we demonstrate that bacteria with CRISPR-Cas resistance are still partially immune to Acr-encoding phage. As a consequence, Acr-phages often need to cooperate in order to overcome CRISPR resistance, with a first phage taking down the host CRISPR-Cas immune system to allow a second Acr-phage to successfully replicate. This cooperation leads to epidemiological tipping points in which the initial density of Acr-phage tips the balance from phage extinction to a phage epidemic. Furthermore, both higher levels of CRISPR-Cas immunity and weaker Acr activities shift the tipping points towards higher phage densities. Collectively these data help to understand how interactions between phage-encoded immune suppressors and the CRISPR systems they target shape bacteria-phage population dynamics.

**Highlights:**

- Bacteria with CRISPR immunity remain partially resistant to Acr-phage
- Sequentially infecting Acr phages cooperate to overcome CRISPR resistance
- Acr-phage epidemiology depends on the initial phage density
- CRISPR resistant bacteria can drive Acr phages extinct

**eTOC blurb:** Some phages encode Acr proteins that block bacterial CRISPR-Cas immune systems. Although CRISPR-Cas can clear the first infection, this Acr-phage still suppresses the host immune system, which can be exploited by other Acr-phages. This is critical for Acr-phage amplification, but this “cooperation” only works beyond a critical Acr-phage density threshold.

## Introduction

Bacteria evolve CRISPR-Cas immunity against bacteriophage (phage) by inserting phage-derived sequences into CRISPR loci on the host genome (Barrangou et al. 2007). Processed transcripts of CRISPR loci guide CRISPR-associated (Cas) surveillance complexes and effector nucleases to detect and destroy complementary genomes of re-infecting phages (Brouns et al. 2008; Garneau et al. 2010). It has been demonstrated that evolution of CRISPR immunity in bacterial populations can drive rapid phage extinction (van Houte et al. 2016). In the face of this strong selection pressure, many phages have evolved to encode anti-CRISPR (*acr*) genes (Bondy-Denomy et al. 2013; Pawluk et al. 2014; Pawluk et al. 2016a), which antagonize CRISPR immune systems of their bacterial hosts by inhibiting CRISPR surveillance complexes or effector nucleases (Bondy-Denomy et al. 2015; Pawluk et al. 2016b; Wang et al. 2016; Wang et al. 2016; Chowdhury et al. 2017; Dong et al. 2017; Guo et al. 2017; Harrington et al. 2017; Hynes et al. 2017; Pawluk et al. 2017; Peng et al. 2017; Rauch et al. 2017; Shin et al. 2017; Yang et al. 2017; Hong et al. 2018). These *acr* genes were first identified in temperate *Pseudomonas* phages (Bondy-Denomy et al. 2013) and can rescue phage from CRISPR-mediated extinction (van Houte et al. 2016). However, previously reported data suggests that their ability to block CRISPR resistance is imperfect, and that some Acrs are more potent than others (Bondy-Denomy et al. 2013). For example, phage encoding AcrF1 had greater levels of infectivity on CRISPR resistant hosts compared to phage encoding AcrF4, but in all cases Acr-phage infectivity was highest on hosts lacking CRISPR-Cas immunity (Bondy-Denomy et al. 2013). While these data suggest that CRISPR immunity provides partial resistance against Acr-phage infection, it has remained unclear how these patterns of partial resistance impact the ability of Acr-phage to replicate and amplify. Here we demonstrate that Acr-phages need to cooperate in order to overcome partial resistance of CRISPR immune hosts. This requirement for cooperation has important epidemiological consequences as it causes Acr-phages to be driven extinct if their initial titers are below a critical threshold value, but allows them to amplify when their titers exceed this threshold.

## Results

### CRISPR-Cas confers partial immunity to Acr-phages

To investigate the consequences of the previously observed partial resistance of CRISPR immune bacteria against Acr-phages, we expressed AcrF1 (from phage JBD30) and AcrF4 (from phage JBD26) in an isogenic phage DMS3*mvir* background, which lacks an endogenous AcrF but is closely related to both parental phages. Consistent with previous observations (Bondy-Denomy et al. 2013), EOP assays with DMS*mvir*-AcrF1 and DMS3*mvir*-AcrF4 confirmed partial immunity of CRISPR-resistant hosts to these Acr-phages and demonstrated that Acrs differ in their ability to block CRISPR resistance, with AcrF1 being a more potent suppressor of CRISPR resistance than AcrF4 (**Fig. 1a**). As expected, EOPs of Acr-phages on WT hosts were higher compared to ancestral phage DMS3*mvir*, which is *a priori* targeted by one spacer of the WT PA14 CRISPR-Cas system, but lower than those of phage DMS3*vir*, which is not *a priori* targeted by the WT CRISPR-Cas system (**Fig. 1a**). Furthermore, EOPs decreased when hosts carried two or five (hereafter named “BIM2” and “BIM5”, Bacteriophage Insensitive Mutant) targeting spacers, presumably because this increases the proportion of surveillance complexes that target the phage (in addition to the targeting spacers, all bacteria encode 35 non-targeting spacers). Furthermore, competition between CRISPR-resistant and sensitive bacteria showed that in the presence of Acr-phages CRISPR resistance provides a fitness advantage (**Fig. 1b**; F_1,53_ = 193.98, p<0.0001), which is consistent with the observation that targeting spacers provide partial resistance to Acr-phage.

**Figure 1.**
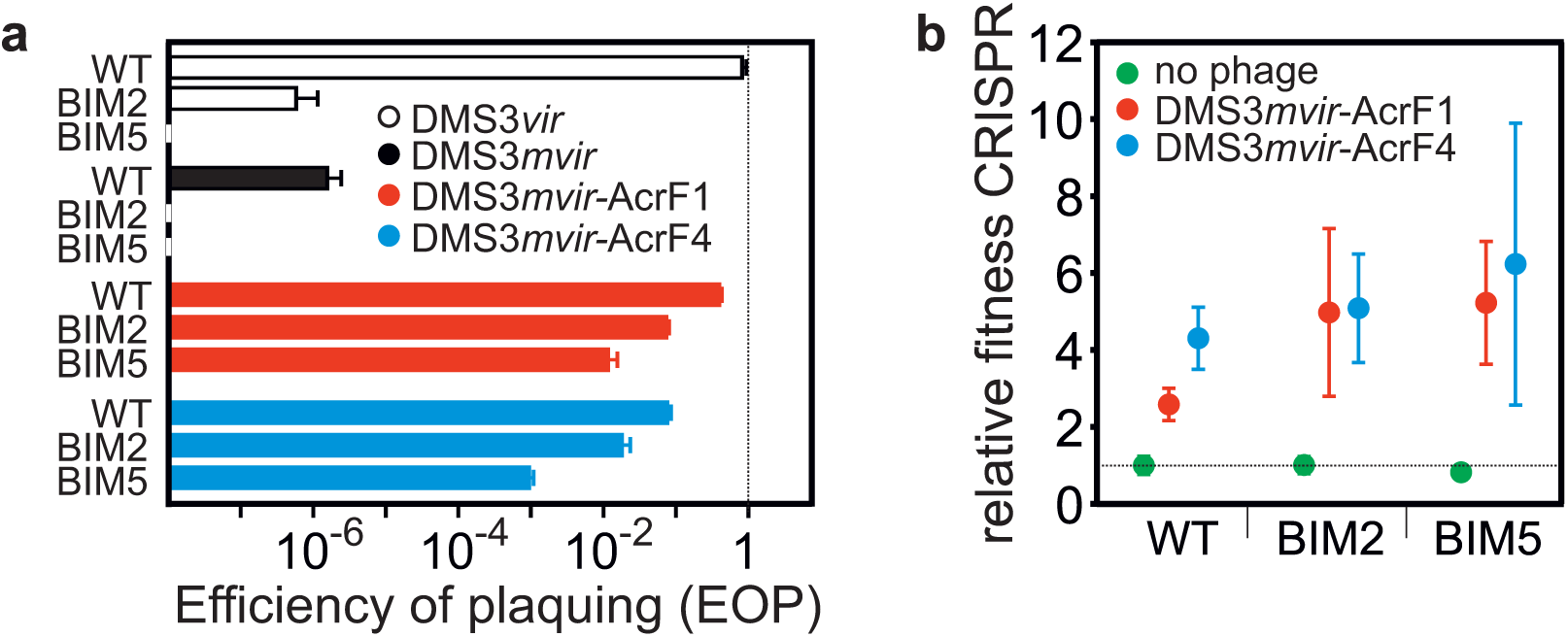
CRISPR-resistant hosts are partially immune to phage encoding an anti-CRISPR**. (a)** Efficiency of plaquing (EOP) of DMS3*vir* (white bars) on PA14 WT (completely sensitive to DMS3*vir*), BIM2 (1 spacer targeting DMS3*vir*) and BIM5 (4 spacers targeting DMS3*vir*); EOP of DMS3*mvir* (black bars), DMS3*mvir*-AcrF1 (red bars) and DMS3*mvir*-AcrF4 (blue bars) on PA14 WT (1 spacer targeting DMS3*mvir*, DMS3*mvir*-AcrF1 and DMS3*mvir*-AcrF4), BIM2 (2 spacers targeting DMS3*mvir*, DMS3*mvir*-AcrF1 and DMS3*mvir*-AcrF4) and BIM5 (5 spacers targeting DMS3*mvir*, DMS3*mvir*-AcrF1 and DMS3*mvir*-AcrF4). Each experiment was performed as 6 independent replicates. Error bars represent 95% confidence intervals (c.i.). **(b)** Relative fitness of CRISPR-resistant bacteria on PA14 WT, BIM2 and BIM5 in the absence of phage (green data points) or in the presence of phage DMS3*mvir*-AcrF1 (red data points) or phage DMS3*mvir*-AcrF4 (blue data points). Each experiment was performed as 6 independent replicates, error bars represent 95% c.i.

### The size of the initial Acr-phage inoculum determines the epidemiological outcome

While full CRISPR resistance can drive phages extinct (van Houte et al. 2016), the phage epidemiology associated with partial resistance to Acr-phages is unclear. We explored this by measuring phage amplification following infection of CRISPR-resistant hosts. Whereas phage always reached similar titers when amplified for 24 hours on CRISPR-KO hosts, independent of the initial phage amount (**Fig. 2a-c**), phage amplification on WT bacteria (one targeting spacer) was dependent on the initial phage amount, with phage DMS3*mvir* amplifying exclusively beyond a threshold of around 10^6^ pfus (**Fig. 2d-f**). For the Acr-phages this effect was even stronger on BIM2 (two targeting spacers) and BIM5 (5 targeting spacers) hosts, revealing epidemiological tipping points that depend both on the level of host resistance and the strength of the Acr (**Fig. 2g-l**). DMS3*mvir*-AcrF1 could only cause an epidemic on BIM2 if the initial amount of phage exceeded a threshold of ∼10^5^ pfus, and was driven extinct below this threshold (**Fig. 2h**), and for DMS3*mvir*-AcrF4 approximately 100-fold more phage was necessary to cause an epidemic (**Fig. 2i**). On BIM5 the tipping point shifted to approximately 10-fold higher phage titers for both Acr-phages (**Fig. 2k,l**).

**Figure 2.**
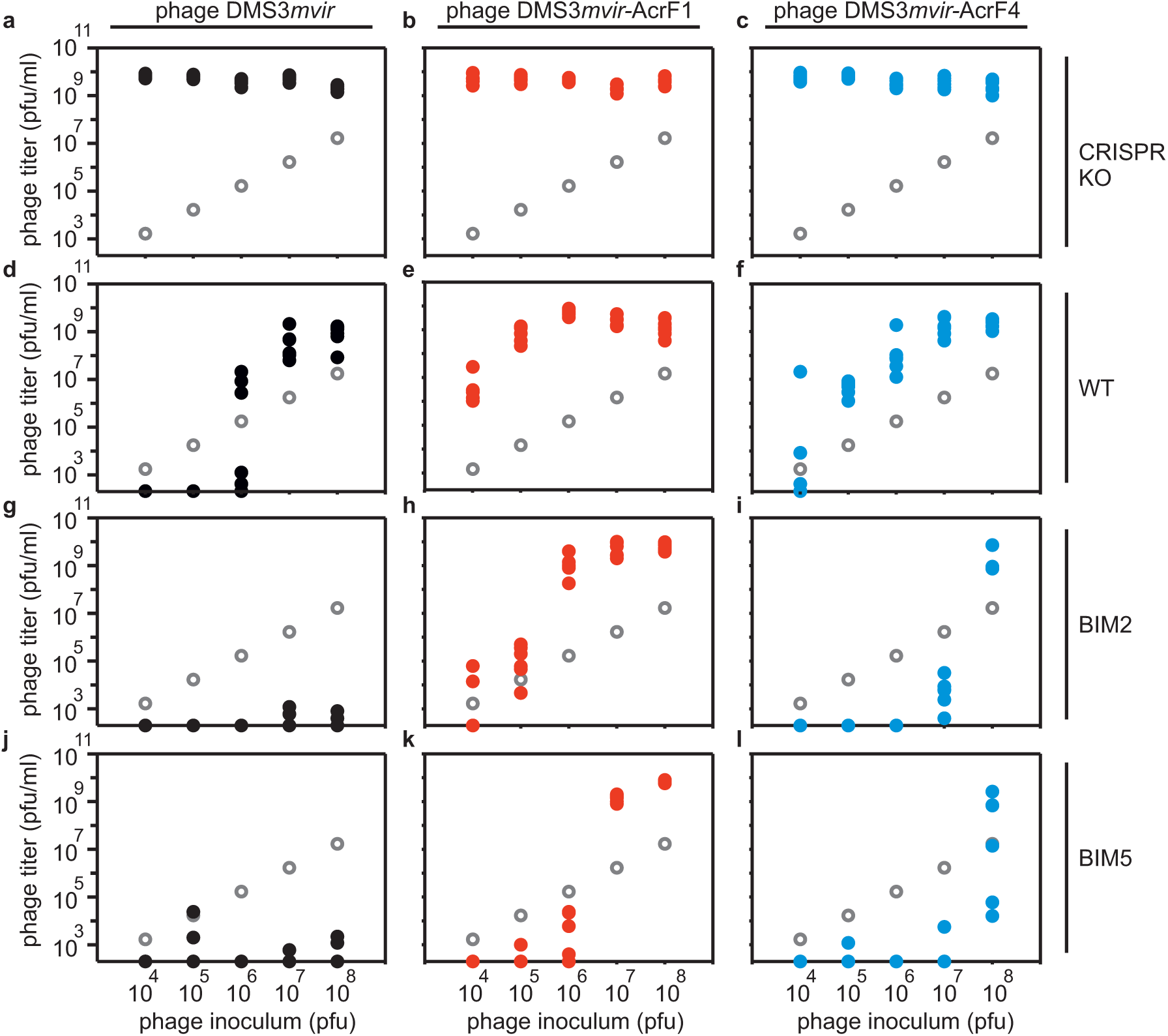
Partial immunity to phages encoding anti-CRISPR causes epidemiological tipping points that are phage density-dependent. Viral titers at 24 hours post-infection (hpi) with DMS3*mvir* (**a,d,g,j**), DMS3*mvir*-AcrF1 (**b,e,h,k**) or DMS3*mvir*-AcrF4 (**c,f,i,l**) of PA14 CRISPR-KO (**a-c**), WT (**d-f**), BIM2 (**g-i**) or BIM5 (**j-l**). Grey circles indicate the phage titers (pfu/ml) at the start of the experiment (corresponding to the addition of 10^4^, 10^5^, 10^6^, 10^7^ or 10^8^ pfus). Colored data points represent phage titers at 24 hpi; each data point represents an independent biological replicate (*n*=6). The limit of detection is 200 pfu/ml.

### Epidemiological tipping points in Acr-phage amplification are not due to phage evolution

Given that phage epidemics only occurred when bacterial cultures were infected with higher amounts of phage, we hypothesized they might be caused by rare phage mutants that “escape” (partial) CRISPR resistance due to mutations in their target sequence (“protospacer”) (Antia et al. 2003; Deveau et al. 2008; Semenova et al. 2011; Jiang et al. 2013; van Houte et al. 2016). To test this, we sequenced phage that was isolated from the observed epidemics on WT, BIM2 and BIM5 following infection with 10^8^ pfus (i.e. from Fig. 2d-f,h,i,k,l). This showed that the epidemic caused by control phage DMS3*mvir* on WT bacteria was indeed caused by phage that carried a mutated protospacer (i.e. mutation in the seed and PAM region) (**Fig. S1a**). However, in the context of Acr-phage, we found only one example, namely that of DMS3*mvir*-AcrF4 on PA14 WT, where the epidemic was associated with a protospacer mutation (**Fig. S1a**). For all other Acr-phage epidemics, protospacer SNP frequencies were similar to those of the ancestral phage (**Fig. S1a**). In these cases we could also not detect any differences in the ability of evolved and ancestral phages to amplify on the CRISPR-resistant hosts they were isolated from **(Fig. S1b)**. Therefore, unless the Acr is weak and the host carries only one spacer, phage evolution cannot explain the observed epidemiological tipping points of Acr-phages.

### Acr-phage amplification is density-dependent

Having ruled out that the observed tipping points by Acr-phage are the result of escape phage evolution, we hypothesized that the density of Acr-phage may determine the observed tipping points. To test this hypothesis, we examined whether amplification of Acr-phage was density-dependent without altering the initial amount of phage. This was done by measuring amplification of the same initial amount of phage on different volumes of bacterial host culture, generating a High Phage-Density (HPD) condition (small volume), and a Low Phage-Density (LPD) condition (large volume). Phage amplification was greater on CRISPR-KO hosts under LPD conditions compared to HPD conditions, simply because the bacterial densities are constant across the treatments and the large volume therefore contains proportionally more bacteria on which the phage can replicate (**Fig. 3**; F_1,22_ = 10.54, p < 0.01). However, when Acr-phages were amplified on CRISPR-resistant hosts (BIM2), the greatest level of amplification was observed under HPD conditions (**Fig. 3**; F_1,22_ = 59.68, p < 0.0001), demonstrating that Acr-phage amplification is indeed positively density-dependent. Furthermore, the level of amplification of Acr-phages on this CRISPR-resistant host was independent of the presence of high amounts of phage DMS3*mvir*, which lacks an *acrF* gene, demonstrating that the observed density-dependence is specifically linked to the density of *acr* genes, and cannot be explained by saturation of CRISPR-Cas complexes with targeted phage genomes (**Fig. S2**).

**Figure 3.**
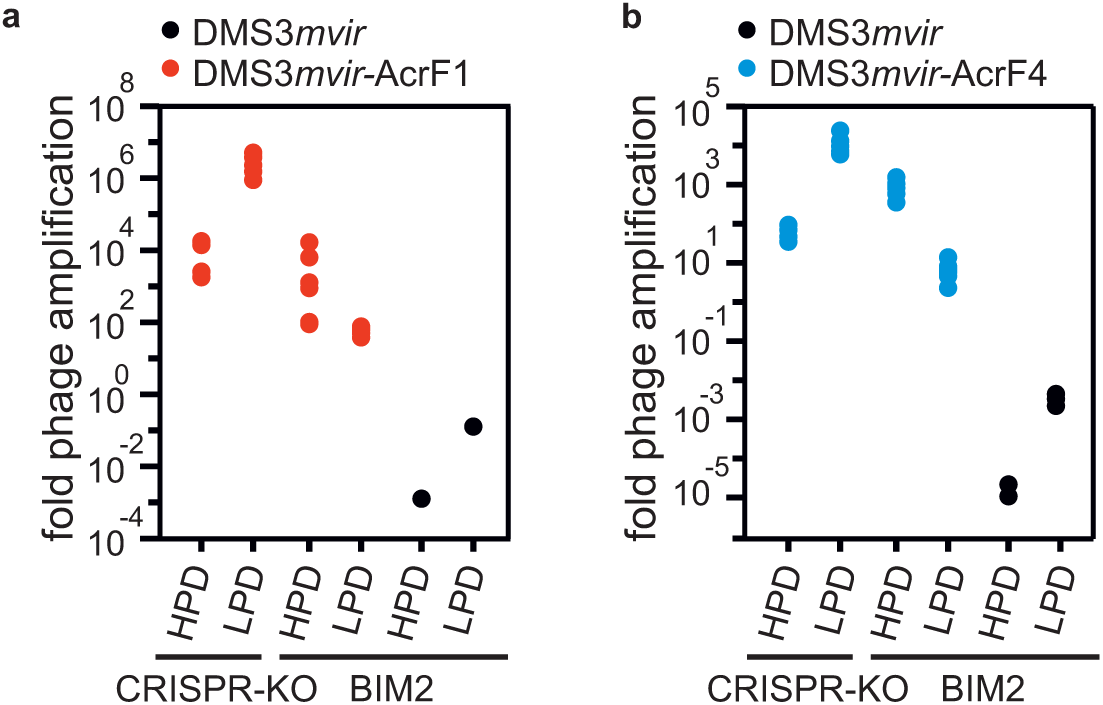
Acr-phage amplification is density-dependent. (**a**) Fold phage amplification at 24 hpi with 10^6^ pfus DMS3*mvir* (black data points) or DMS3*mvir*-AcrF1 (red data points) of PA14 CRISPR-KO (sensitive) or BIM2 under either high phage densities (HPD, 6 ml culture) or low phage densities (LPD, 600 ml culture). (**b**) Fold phage amplification at 24 hpi with 10^8^ pfus DMS3*mvir* (black data points) or DMS3*mvir*-AcrF4 (blue data points) of PA14 CRISPR-KO (sensitive) or BIM2 under either HPD or LPD; each data point represents an independent biological replicate (*n*=6). The limit of detection is 200 pfu/ml.

### Epidemiological tipping points can result from cooperation between sequentially infecting Acr-phages

The observed density-dependent phage amplification suggested that Acr-phages may cooperate in order to successfully amplify. For example, if co-infections were required to effectively suppress host resistance, epidemiological tipping points could correspond to parasite densities where co-infections become common (Regoes et al. 2002). However, this hypothesis is unlikely to explain our results, because the tipping points occurred at multiplicity of infection (MOI) values where co-infections are expected to be rare (e.g. MOI∼0.06 for DMS3*mvir*-AcrF1 on BIM2). To explore what factors may cause the observed epidemiological tipping points at low MOIs and in the absence of phage evolution, we generated a theoretical model (**Supplemental Data**) that accounts for the efficacy *ρ* of CRISPR resistance in the bacteria (*ρ* increases with the number of spacers targeting the phage) as well as the efficacy ϕ of Acr in the phage (consistent with the EOP data). In this form, the model predicts that upon infection of 10^6^ CRISPR-resistant bacteria phage expressing a strong Acr can always amplify, regardless of the initial phage density (**Fig. S3**, *ϕ*=0.67, purple line), whereas phage with a weak Acr can never amplify (**Fig. S3**, *ϕ*=0.6 and *ϕ*=0.5, magenta and green lines respectively; grey lines correspond to the initial amount of phage and values below this line indicate a lack of phage amplification). Given that these model predictions are inconsistent with our experimental data, we then extended the model by incorporating the assumption that during failed infections some Acr is produced that causes the surviving host to enter a “suppressed” state (“S”). This immunosuppression decreases the efficacy of host resistance and allows following phages to exploit these bacteria (**Fig. 4a**). Crucially, if the immunosuppressed state is assumed, the model predicts epidemiological tipping points and, in accordance with our empirical data, these tipping points occur at MOIs far below 1 (**Fig. 4b**). Besides, our experimental observations that the position of the tipping points shifts when Acrs are weaker or host resistance is stronger (**Fig. 2**) are fully explained by our model when we vary the effect of the efficacy ϕ of the Acr in the phage (**Fig. 4b**), or the efficacy *ρ* of CRISPR resistance in the bacteria (i.e., the number of spacers in the host targeting the phage, **Fig. 4c**). Moreover, longer periods of immunosuppression shift the tipping points to lower phage densities, as it increases the probability that a host will be re-infected when it is still in the immunosuppressed state (**Fig. 4d,** indicated by *γ*). This model therefore predicts that Acr-phage infections can cause CRISPR-resistant bacteria to become immunosuppressed, allowing cooperation between sequentially infecting Acr-phages to overcome CRISPR immunity, which is a critical factor in determining whether Acr-phages can amplify.

**Figure 4.**
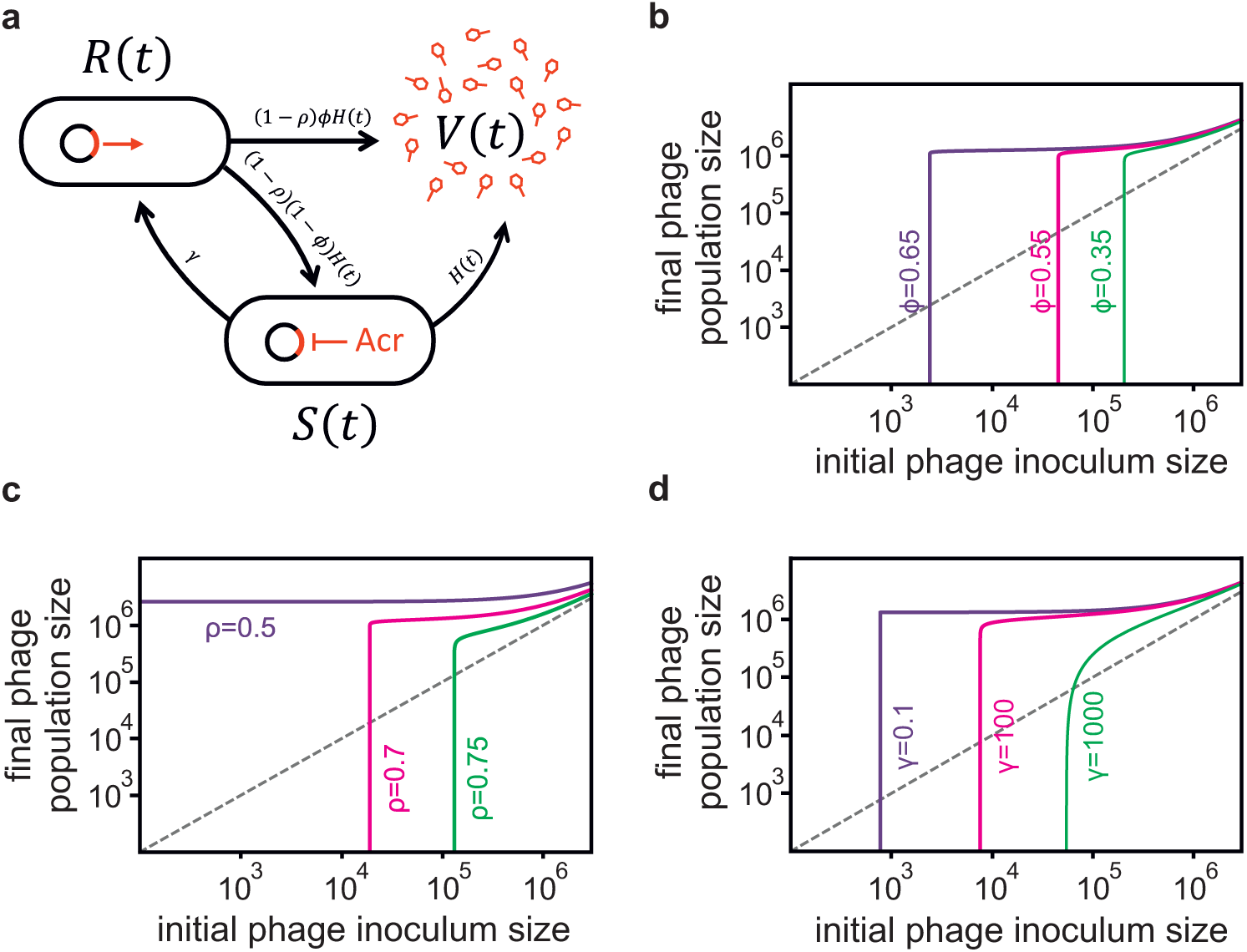
Lasting immunosuppression following unsuccessful Acr-phage infections can lead to epidemiological tipping points. (**a**) Infection model of the Acr-phage (see details of the model in Supplementary Information). The parameter *H(t)*=*αV(t)* refers to the rate at which bacteria are infected by free phage particles. (**b**) Effect of initial Acr-phage inoculum density on the phage density at 24 hpi for different values of Acr efficacy (*ϕ* = 0.65 (purple), *ϕ* = 0.55 (magenta) and *ϕ* = 0.35 (green)); other parameter values: *B* = 5, *α* = 0.001, *ρ* = 0.7, *γ* = 20. (**c**) Effect of initial Acr-phage inoculum density on the phage density after 24h of incubation for different values of CRISPR efficacy (*ρ* = 0.5, 0.7 and 0.75; purple, magenta, green respectively); other parameter values: *B* = 5, *α* = 0.001, *ϕ* = 0.6, *γ* = 20. (**d**) Effect of initial Acr-phage inoculum density on the phage density after 24h of incubation for different values of the duration of the immunosuppressive state (*γ* = 0.1, 100 and 1000; purple, magenta, green respectively); other parameter values: *B* = 5, *α* = 0.001, *ϕ* = 0.65, *ρ* = 0.7. In all graphs, grey lines correspond to the initial amount of phage and values below this line indicate a lack of phage amplification.

### Unsuccessful infections by Acr-phages cause CRISPR-resistant hosts to become immunosuppressed

To validate this model we tested the key assumption that unsuccessful infections by Acr-phages cause CRISPR-resistant hosts to become immunosuppressed. To this end, we pre-infected BIM2 and BIM5 bacteria with Acr-phage at a low MOI (∼0.3) and subsequently washed away all remaining phages from the culture. We then measured the relative transformation efficiency (RTE) of the surviving cells by transforming pre-infected bacteria with either a CRISPR-targeted plasmid (T) or a non-targeted plasmid (NT). For all phage treatments, the RTE of pre-infected CRISPR-KO bacteria and non-infected controls were not significantly different, as expected (**Fig. 5**; F_1,51_ = 1.12, p = 0.35). However, when BIM2 or BIM5 bacteria were pre-infected with phage DMS3*mvir*-AcrF1, the RTE increased significantly compared to the DMS3*mvir* and no-phage controls (**Fig. 5**; BIM2: F_1,17_ = 26.82, p < 0.0001, BIM5: F_1,20_ = 20.16, p < 0.0001), demonstrating lasting immunosuppression of CRISPR-resistant hosts following an unsuccessful infection with Acr-phage. Consistent with its weaker Acr activity, lasting immunosuppression following infection with DMS3*mvir*-AcrF4 was only observed in BIM2 (**Fig. 5**; F_1,17_ = 5.26, p<0.05) and not BIM5 (F_1,20_ = 2.07, p = 0.15).

**Figure 5.**
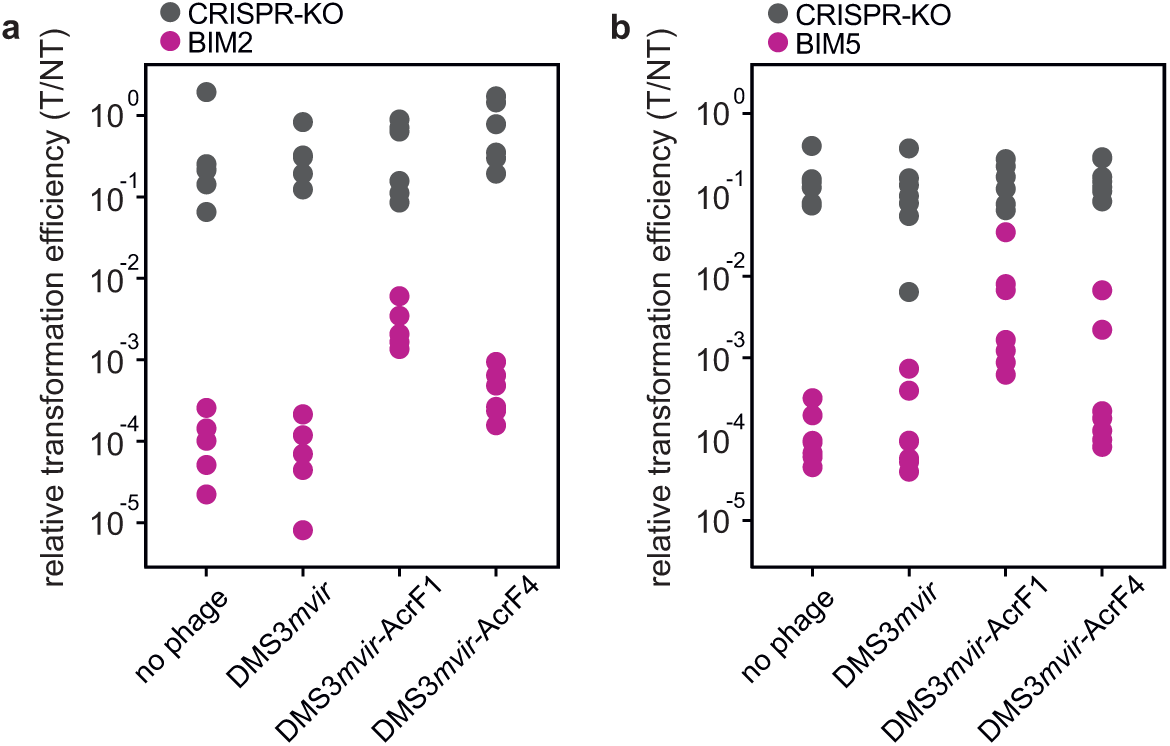
Infection with Acr-phage lead to long-term immunosuppression of CRISPR-resistant hosts. (**a**) Relative transformation efficiencies (RTE) of CRISPR-KO (grey data points) or BIM2 (purple data points) pre-infected with 1.6*10^9^ pfus of either DMS3*mvir*, DMS3*mvir*-AcrF1 or DMS3*mvir*-AcrF4, or not phage-infected. Each data point represents an independent biological replicate (*n*=6). (**b**) RTE of CRISPR-KO (grey data points) or BIM5 (purple data points) pre-infected as described for (a). Each data point represents an independent biological replicate (*n*=7).

## Discussion

The discovery of Acr proteins has been a major breakthrough in CRISPR-Cas research (Bondy-Denomy et al. 2013). Since their initial discovery much progress has been made towards biochemical characterization of Acrs and the unraveling of their molecular mode of action. Here we study the population dynamics associated with CRISPR-Acr interactions and demonstrate that the initial density of Acr-phages that infect bacteria with partial CRISPR resistance determines whether phages go extinct or amplify. We ruled out that the need for high Acr-phage densities is simply linked to the evolution of phage escape mutations. Instead, the observed epidemiological tipping points can be primarily explained by long-term suppression of CRISPR resistance following an unsuccessful infection, which is consistent with the slow dissociation kinetics of Acr-Cas protein complexes (Chowdhury et al. 2017). Modeling the temporal phage and host population dynamics shows that during the initial stages of infection Acr-phage densities decline due to the high proportion of unsuccessful infections (Supplemental Information and **Fig. S4ab**). However, as the densities of immunosuppressed hosts increase a greater proportion of infections becomes successful. If the initial densities of Acr-phages are high enough, densities of immunosuppressed hosts reach a critical threshold where the amount of new Acr-phages that are produced from successful infections outweighs the loss of Acr-phages due to unsuccessful infections, causing the epidemic to take off (**Fig. S4b**). If this critical threshold is not reached, the Acr-phage goes extinct and immunosuppressed hosts revert to their resistant state (**Fig. S4a**).

The long-term immunosuppression following a failed infection of Acr-phage is cooperative in that it provides a benefit to Acr-phages that sequentially infect the same host to overcome CRISPR resistance. Future studies aimed at measuring the costs and benefits associated with phage-encoded Acr genes and their natural ecology will be critical to understanding the evolutionary drivers and stability of Acr-phage cooperation (Sachs et al. 2004, West et al 2007). Specifically, it is unclear the extent to which lasting immunosuppression evolved because of the indirect fitness benefits associated with the enhanced infection success of clone mates in the population, or whether it is primarily a by-product of the direct individual-level benefits of suppressing the host immune system.

As discussed above, amplification of Acr-phages on CRISPR-resistant bacteria was found to occur only if the initial Acr-phage densities were above a certain threshold value, which is determined both by the strength of the Acr and the resistance level of the host. Below this threshold CRISPR-resistant bacteria drive phage to extinction, despite the phage-encoded Acr. In general, positive density-dependent fitness effects are thought to play an important role in various ecological contexts, such as species invasions, extinctions and disease epidemics (Courchamp et al. 1999; Stephens et al. 1999). Existing theory predicts that parasite density-dependent tipping points in disease epidemics can occur when the infection dynamics of an individual host depends on the parasite dose, for example when there is a threshold in the number of co-infecting parasites that are required to establish a successful infection (Regoes et al. 2002). This work shows that epidemiological tipping points can also take place under conditions where parasite densities are too low for co-infections to be common if unsuccessful infections leave behind an immunosuppressed host. The profound epidemiological consequences that were found to be associated with lasting immunosuppression in our empirical system warrant future studies to explore whether similar effects play a role in the epidemiology of other infectious diseases.

## Acknowledgements

ML was supported by funding from the Wellcome Trust (https://wellcome.ac.uk) (109776/Z/15/Z), which was awarded to ERW. ERW further acknowledges the Natural Environment Research Council (http://www.nerc.ac.uk) (NE/M018350/1), the BBSRC (BB/N017412/1) and the European Research Council (https://erc.europa.eu) (ERC-STG-2016-714478 - EVOIMMECH) for funding. SVH acknowledges funding from the People Programme (Marie Curie Actions; https://ec.europa.eu/research/mariecurieactions/) of the European Union‘s Horizon 2020 (REA grant agreement no. 660039) and from the BBSRC (BB/R010781/1). The authors thank Adair Borges and Joe Bondy-Denomy (UCSF) for providing the DMS3*mvir*-AcrF4 recombinant phage. Anne Chevallereau is acknowledged for providing feedback on the manuscript.

## Author contributions

Conceptualization, ERW, SvH, AB; Methodology, ERW, SvH, ML, SG; Investigation, ML, HC, SvH, ERW; Formal Analysis, ML, SM, SvH, ERW, SG; Writing – Original Draft, SvH, ERW; Writing – Review & Editing, SvH, ERW, ML, SG, AB; Funding Acquisition, ERW, SvH; Supervision, ERW, SvH.

## Declaration of interest

The authors declare no competing interests.

## Materials and methods

### Bacterial strains

*P. aeruginosa* UCBPP-PA14 (WT; carrying one spacer targeting DMS3*mvir*) and *P. aeruginosa* UCBPP-PA14 *csy3::LacZ* (referred to as CRISPR-KO, which carries a disruption of an essential *cas* gene and can therefore not evolve CRISPR immunity), have been described in (Cady et al. 2012; Westra et al. 2015; van Houte et al. 2016). The BIM2 strain (carrying two spacers targeting DMS3*mvir*) has been described in (Westra et al. 2015). The BIM5 strain (carrying 5 spacers targeting DMS3*mvir*) was generated by challenging PA14 BIM2 bacteria with escape phage in multiple rounds, giving rise to BIM3, BIM4 and finally BIM5.

### Phage strains and generation of recombinant phages

Phages DMS3*vir*, DMS3*mvir* and DMS3*mvir*-AcrF1 have been previously described in refs. (Cady et al. 2012; Bondy-Denomy et al. 2013; van Houte et al. 2016). Phage DMS3*mvir*-AcrF4, which carries the anti-CRISPR gene *acrF4* (formerly JBD26-37), was made by generating a plasmid possessing the *acrF4* gene flanked by regions of homology from JBD30 (which are nearly identical to DMS3*m*). DMS3*m* was used to infect cells possessing this plasmid and recombinants possessing *acrF4* were selected for. This phage (DMS3*m*-AcrF4) was then made virulent via C-repressor deletion as in (Cady et al. 2012). Genome assemblies of DMS3*mvir*-AcrF1 and DMS3*mvir*-AcrF4 used in this study can be found under GenBank accession number XXXXX.

### Efficiency of Plaquing (EOP) assays

EOP assays were carried out on plates containing LB with 1.5% agar. A mixture of molten soft LB agar (0.5%), 300 μl of bacteria (grown O/N in M9 growth medium supplemented with 0.2% glucose) and 100 μl of serially diluted phage was poured on top of the hard agar layer. Plates were incubated O/N at 37 ºC and plaques were enumerated the next day.

### Competition assays to measure CRISPR-associated fitness

Competition experiments were performed in glass vials in 6 ml M9 medium (22 mM Na_2_HPO_4_; 22 mM KH_2_PO_4_; 8.6 mM NaCl; 20 mM NH_4_Cl; 1 mM MgSO_4_; 0.1 mM CaCl_2_) supplemented with 0.2% glucose. Competition experiments were initiated by inoculating 1:100 from a 1:1 mixture of overnight cultures (grown in M9 medium + 0.2% glucose) of the CRISPR-resistant strain (either WT, BIM2 or BIM5) and the sensitive CRISPR-KO strain. To each microcosm 10^4^ plaque forming units (pfus) of either DMS3*mvir*-AcrF1 or DMS3*mvir*-AcrF4 phage was added, and each treatment was performed in six independent replicates. At 0 and 24 hours after the start of the competition experiment samples were taken and cells were serially diluted in M9 salts (22 mM Na_2_HPO_4_; 22 mM KH_2_PO_4_; 8.6 mM NaCl; 20 mM NH_4_Cl) and plated on LB agar supplemented with 50 μg.ml^-1^ X-gal (to allow discrimination between CRISPR-resistant (white) and sensitive CRISPR-KO (blue) bacteria). For all competitions, a control competition experiment was performed in the absence of phage. For all competitions, relative frequencies of the CRISPR-resistant strains were determined and used to calculate the relative fitness (rel. fitness = [(fraction strain A at t=x) * (1 – (fraction strain A at t=0))] / [(fraction strain A at t=0) * (1 – (fraction strain A at t=x)]). These values were used for student t-test or ANOVA. All subsequent statistical analyses were carried out using R.

### Infection assays in liquid medium

Infection assays were performed in glass vials by inoculating 6 ml M9 medium supplemented with 0.2% glucose with 60 μl (app. 2*10^7^ colony forming units (cfus) bacteria from fresh overnight cultures (also grown in M9 medium + 0.2% glucose) of either the CRISPR-KO, WT, BIM2 or BIM5 strain. To these microcosms 10^4^, 10^5^, 10^6^, 10^7^ or 10^8^ pfus of either DMS3*mvir*, DMS3*mvir*-AcrF1 or DMS3*mvir*-AcrF4 phage were added. Each treatment was performed in six independent replicates. Microcosms were incubated at 37 ºC while shaking at 180 rpm. Phage was extracted at 24 hours after the start of the experiment by chloroform extraction on all samples (sample: chloroform 10:1 v/v), and phage titers were determined by spotting isolated phage samples on a lawn of CRISPR-KO bacteria.

For the experiment shown in **Fig. S2** the same protocol was used but with 10^8^ pfus DMS3*mvir* added to each treatment at the start of the experiment (**Fig. S2b,d**). For the infection assays in which phage densities were manipulated by using different volumes of growth medium (**Fig. 3**), glass microcosms containing 6 ml M9 growth medium + 0.2% glucose were used for the high phage-density (HPD) treatment. These were inoculated with 60 μl (app. 2*10^7^ cfus) bacteria from fresh overnight cultures of the CRISPR-KO or BIM2 strains, as above. For the low phage-density (LPD) treatment Duran 1L glass bottles containing 600 ml M9 growth medium + 0.2% glucose were used, which were inoculated with 6 ml (app. 2*10^9^ cfus) bacteria from the same overnight cultures. To both glass vials and bottles phage was added (10^6^ pfus of DMS3*mvir*-AcrF1; 10^8^ pfus of DMS3*mvir*-AcrF4 or 10^8^ pfus of DMS3*mvir*). Glass vials and bottles were incubated at 37 ºC while shaking at 180 rpm. Total phage was extracted at 24 hours after the start of the experiment by chloroform extraction (sample: chloroform 10:1 v/v), and phage titers were determined by spotting all isolated phage samples on a lawn of CRISPR-KO bacteria. All experiments were performed in six independent replicates.

### Deep sequencing of phages

From the experiments shown in Fig. 2a-l, phage was isolated from the treatments that were infected with 10^8^ pfus of phage. These phages were used for a new round of infection assays in liquid media (Fig. S1), effectively following the methods described above, and for deep sequencing analysis. To obtain sufficient material for the latter, isolated phage was amplified by plaque assay on the CRISPR-KO strain. Phage samples from all replicates within a single treatment were pooled. As controls, ancestral DMS3*mvir*, DMS3*mvir*-AcrF1 and DMS3*mvir*-AcrF4 phage were processed in parallel. Phage genomic DNA extraction was performed with 600 μl sample at approximately 1012 pfu/ml using the Norgen phage DNA isolation kit, following the manufacturer‘s instructions. Barcoded Illumina Truseq Nano libraries were constructed from each DNA sample with an approximately 350 bp insert size and 2 × 250 bp reads generated on an Illumina MiSeq platform. Reads were trimmed using Cutadapt version 1.2.1 and Sickle version 1.200 and then overlapping reads merged using Flash version 1.2.11 to create high quality sequence at approximately 8000 × coverage of DMS3*mvir* per sample. These reads were mapped to the DMS3*mvir* genomes using bwa mem version 0.7.12 and allele frequencies of single nucleotide polymorphisms within viral target regions quantified using samtools mpileup version 0.1.19. Further statistical analyses was performed in R version 3.4.1. Sequence data are available from the European Nucleotide Archive under accession number PRJEB25016 and analysis scripts are available from https://github.com/s-meaden/landsberger.

### Generation of CRISPR-targeted plasmid

For the long-term immunosuppression experiment (**Fig. 5**) plasmid pHERD30T was used. To generate pHERD30T that was *a priori* targeted by the PA14 WT Type I-F CRISPR-Cas system, the mutant plasmid pHERD30T*targ* was generated that carries a 32-nt protospacer sequence flanked by a GG protospacer adjacent motif (PAM) with full complementarity to spacer 1 in CRISPR locus 2 of the PA14 WT Type I-F CRISPR-Cas system. This was done by ligation of oligonucleotides that upon annealing create overhangs that are compatible with *Hind*III ligation (5’-agcttACCGCGCTCGACTACTACAACGTCCGGCTGATGGa-3’ and 5’-agcttCCATCAGCCGGACGTTGTAGTAGTCGAGCGCGGTa-3’, *Hind*III overhangs in small caps, protospacer sequence in capitals and PAM underlined) in the *Hind*III-digested pHERD30T vector.

### Long-term immunosuppression experiment

Suppression of CRISPR immunity by Acr was measured through a transformation assay. CRISPR-KO, BIM2 or BIM5 bacteria (app. 5*10^9^ cfus) were either not infected or infected with 1.6*10^9^ pfus DMS3*mvir*, DMS3*mvir*-AcrF1 or DMS3*mvir*-AcrF4 in 50 mL Falcon tubes containing 10 mL of Luria-Bertani (LB) broth, and incubated at 37 ºC while shaking at 180 rpm for 2 hours. Bacteria were harvested by spinning at 3500 rpm for 30 minutes, after which a sample was taken from the supernatant for chloroform extraction (sample: chloroform 10:1 v/v) to quantify phage titers (data not shown). The bacterial pellet was then washed twice in 1 mL of a 300 mM sucrose solution to make them competent (Choi et al. 2006). Finally the pellet was resuspended in 300 μl of 300 mM sucrose, and 100 μl from this was used for plating on LB agar to enumerate total bacterial cfus after the infection and sucrose-washing steps (data not shown). The remaining 200 μl was divided in equal volumes over two eppendorf tubes, and were used for electrotransformation with either plasmid pHERD30T (not targeted by CRISPR-Cas; NT) or pHERD30T*targ* (targeted by CRISPR-Cas; T). Electroporated bacteria were resuspended in 1 ml LB broth and incubated 1h at 37ºC at 180 rpm. Bacteria were then pelleted and resuspended in 100 μl LB and plated on LB agar plates containing Gentamycin (50 μg.ml^-1^) and incubated for 16h at 37 ºC to allow transformants to grow. For each of the four phage treatments the number of transformants from the NT and T transformation was used to calculate the relative transformation efficiency (T/NT).

